# A minimal model of T cell avidity may identify subtherapeutic vaccine schedules

**DOI:** 10.1101/2020.12.06.413864

**Authors:** Adarsh Kumbhari, Danya Rose, Peter P. Lee, Peter S. Kim

## Abstract

T cells protect the body from cancer by recognising tumour-associated antigens. Recognising these antigens depends on multiple factors, one of which is T cell avidity, i.e., the total interaction strength between a T cell and a cancer cell. While both high- and low-avidity T cells can kill cancer cells, durable anti-cancer immune responses require the selection of high-avidity T cells. Previous experimentation with anti-cancer vaccines, however, has shown that most vaccines elicit low-avidity T cells. Optimising vaccine schedules may remedy this by preferentially selecting high-avidity T cells. Here, we use mathematical modelling to develop a simple, phenomenological model of avidity selection that may identify vaccine schedules that disproportionately favour low-avidity T cells. We calibrate our model to our prior, more complex model, and then validate it against several experimental data sets. We find that the sensitivity of the model’s parameters change with vaccine dosage, which allows us to use a patient’s data and clinical history to screen for suitable vaccine strategies.

## 1 Introduction

T cells maintain anti-tumour immunity by recognising and killing cancer cells. T cells recognise these cancerous cells through a surface protein—the T cell receptor (TCR)—binding to molecules known as peptide major histocompatibility complexes (pMHCs), which reside on the surface of cancer cells (Murphy, 2011). The overall strength of these TCR-pMHC interactions is termed avidity (Abbas et al., 2014).

Several studies have shown that the selection of high-avidity T cells may be a requirement for durable tumour eradication in certain cancers such as melanoma (Molldrem et al., 2003; Chung et al., 2014). Low-avidity T cells, by contrast, are weakly-tumour killing (Stuge et al., 2004) and may even temper anti-tumour activity by selectively inhibiting high-avidity T cells (Chung et al., 2014). Indeed, experimental evidence suggests that certain cancer vaccines may promote the expansion of low-avidity T cells (Stuge et al., 2004; Rezvani et al., 2011), which may explain why these vaccines cannot maintain durable anti-tumour immunity in clinical trials (Schwartzentruber et al., 2011; Sosman et al., 2008).

To remedy this, multiple techniques have been proposed. These techniques range from searching through peptide libraries to identify peptides that will stimulate high-avidity T cells (McMahan et al., 2006), to harnessing the plasticity of naive T cells to promote their differentiation into high-avidity T cells (Kroger and Alexander-Miller, 2007). More recently, evolutionary principles have been used to select for high-avidity T cells (Bassan et al., 2019). Complementing these experimental studies are mathematical models that aim to improve the efficacy of cancer vaccines, namely treatment schedules (i.e., vaccine dose and timing), from different perspectives. For example, in Sigal et al. (2019), the authors optimise treatment schedules to maximise the clearance of cancer stem cells by killer T cells. Moreover, in Wei et al. (2017) the authors optimise the injection of helper T cells to enhance cytokine-mediated tumour clearance. More broadly, in Joshi et al. (2009), the authors examine how vaccine schedules can be leveraged to avoid tumour recurrence. Besides these studies, researchers have also sought to optimise vaccine schedules in the context of combination therapies. For example, Lai and Friedman (2017) examine how immune checkpoint blockers can be combined with cancer vaccines for enhanced anti-tumour immunity, while Wilson and Levy (2012) look at how a regulatory-protein inhibitor can be combined with a cancer vaccine to induce anti-tumour immunity. Indeed, our own previous modelling work found that vaccine schedules, when optimised, may elicit high-avidity T cells (Kumbhari et al., 2020b,a). Our model, however, is complex, and this complexity makes experimental validation difficult. Moreover, this complexity introduces an element of model uncertainty as not all immune pathways and processes are well understood.

To address this, we develop a simple phenotypic ordinary differential equation (ODE) model that can reproduce the results of our prior model. We validate our model against in vivo murine data from Hailemichael et al. (2013), ex vivo human data from Rezvani et al. (2011) and in vitro data from Wu et al. (2017) and Cawthon et al. (2001). Notably, the model presented here is a reduction of the model developed in Kumbhari et al. (2020a), obtained not via a formal model reduction, but rather via a conceptual reduction informed by our sensitivity analysis from Kumbhari et al. (2020a) and a review of the biological literature. Specifically, our model is based on the experimental observations that (1) mature DCs present antigens at different levels; (2) low DC antigen loads activate only high-avidity T cells, while high DC antigen loads activate both low- and high-avidity T cells; and (3) a history of antigen exposure attenuates T-cell expansion.

We find that the sensitivity of the model’s parameters, which are abstractions of different biological processes, vary with dosage. We use this sensitivity analysis to eliminate inappropriate vaccine schedules (i.e., a schedule that promotes low-avidity T cells) based on a patient’s underlying conditions. This increases the likelihood of electing high-avidity T cells and thus, the likelihood of durable anti-tumour responses. While our study still requires experimental validation, it nevertheless provides a vital proof-of-concept basis for further development of this approach.

## 2 Model

In this section, we develop a minimal model of T cell avidity. Our minimal model establishes a framework for systematically incorporating additional complexity, which may help in quantifying the extent to which different pathways impede tumour clearance. Moreover, in the context of optimising vaccine schedules, our model is amenable to more sophisticated optimisation techniques (that are beyond the scope of this study) such as numerical optimal control. Finally, we note that while no model is perfect, by using only well understood phenotypes of avidity selection, we are able to reduce any model uncertainty in our predictions.

In developing a minimal model of avidity selection, however, we exclude many aspects of the immune response. For example, for example our model does not account for certain cell populations such as natural killer cells, regulatory T cells and helper T cells. We also omit signalling pathways such as cytokine secretion. Importantly, our goal here is to develop a caricature model with a plausible biological basis, rather than a model that aims to capture all known T cell dynamics.

To this end, we assume immature dendritic cells (iDCs) take up antigen and start maturing upon contact with the injected vaccine due to tumour-associated peptides and maturation signals such as vaccine adjuvant, danger signals, or tissue derived immunogenic signals Coffman et al. (2010); Gardner and Ruffell (2016). Maturing DCs migrate to draining lymph nodes, where they present antigens to antigen-specific naive T cells, resulting in their activation to effector T cells (Murphy, 2011; Abbas et al., 2014). Importantly, different DCs present varying levels of antigen on their surfaces, affecting the avidity of T cells that are activated. For simplicity, we focus on the dynamics of killer T cells that are cytolytic against tumours and are the primary target of anti-cancer vaccines (Lollini et al., 2006; Chung et al., 2014; Peng et al., 2019).

To model these interactions, we consider several populations: *P*, the concentration of vaccine peptides; *I*, the concentration of iDCs; *M_L_* and *M_H_*, the concentrations of mature DCs expressing low or high levels of vaccine peptide on their surfaces; and *T_L_* and *T_H_*, the concentrations of killer T cells of low and high avidity. A diagram of the different interactions between these populations is shown in Figure 1. We model the interactions between these populations with an ODE system:

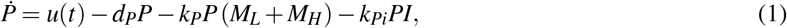

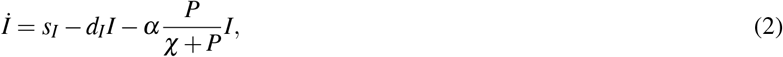

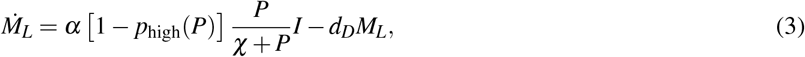

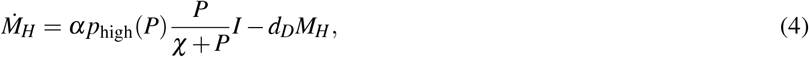

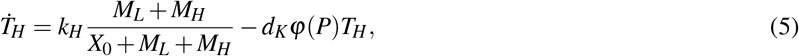

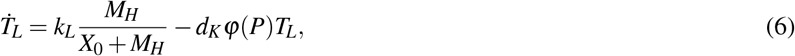

where *u*(*t*) is the vaccine injection rate.

**Fig. 1.**
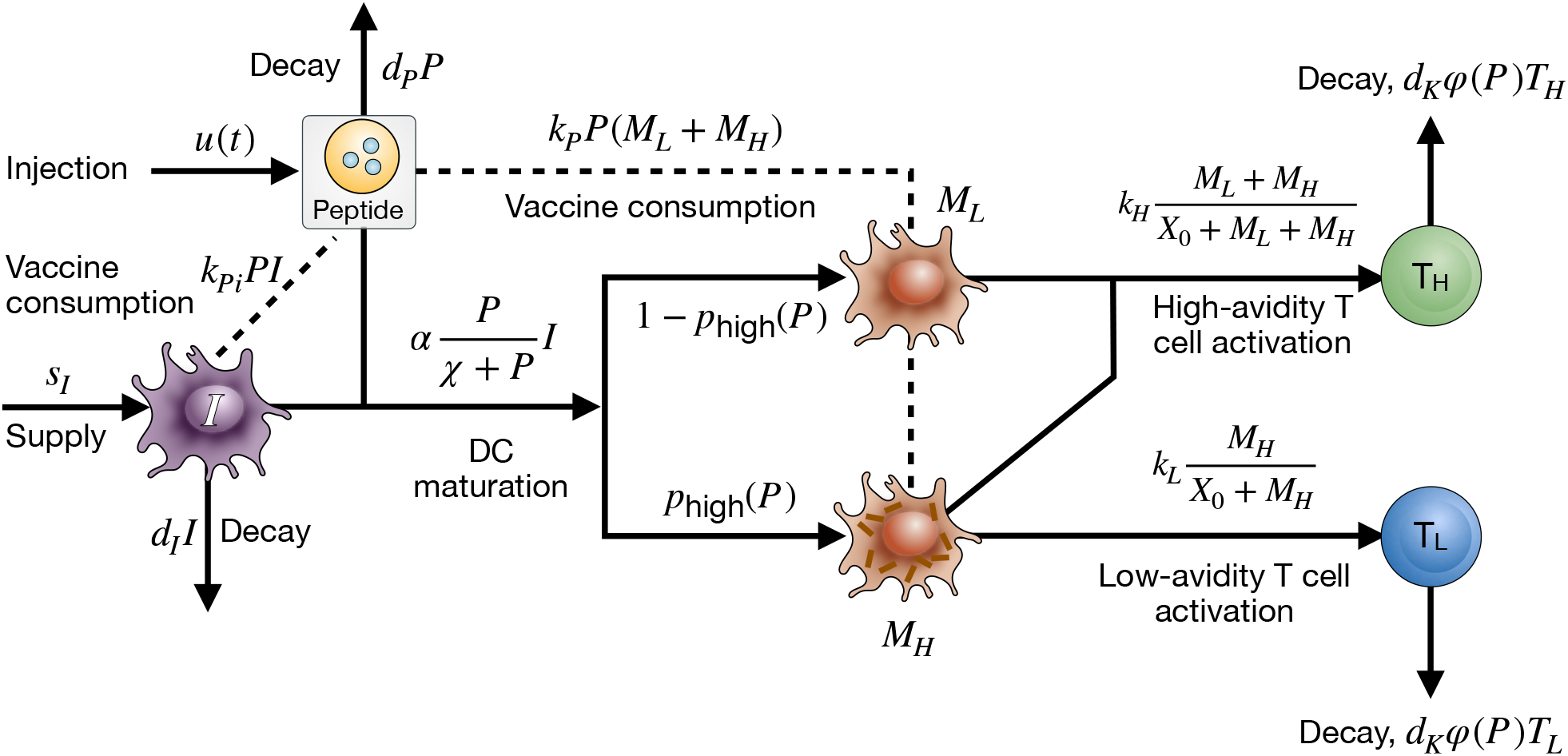
A diagram of the key interactions between injected vaccine peptides, *P*; immature DCs, *I*; mature DCs with different levels of antigen expression, *M_L_* (low) and *M_H_* (high); and a population of low- and high-avidity killer T cells, *T_L_* and *T_H_*

In Eq. (1), vaccine peptides are injected at rate *u*(*t*), decay at rate *d_P_*, and are consumed by mature DCs at rate *k_p_* and by immature DCs are rate *k_Pi_*. In Eq. (2), iDCs are replenished at rate *s_I_* and turnover at rate *d_I_*. The final term in Eq. (2), models the maturation of iDCs due to adjuvant. Because adjuvant is usually not antigen-specific (Garcon and Di Pasquale, 2017), as a simplifying assumption we assume that *all* peptides within the periphery of an iDC are presented at rate *α*. It follows that if the concentration of non-vaccine proteins is denoted by *χ*, then the proportion of peptides presented that are vaccine-associated is *P*/ (*χ* + *P*). Together, these equate to a net flux of *αP*/ (*χ* + *P*).

In Eq. (3) and Eq. (4), iDCs transition into mature DCs at rate 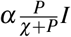 and turnover at rate *d_D_*. The specific probability of transitioning to a mature DC presenting low levels of surface antigens is 1 – *p*_high_(*P*), while the probability of transitioning to a mature DC presenting high levels of surface antigens is *p*_high_(*P*).

Finally, in Eq. (5) and (6), killer T cells activate and proliferate as a function of DC concentration, and decay at rate *d_K_φ*(*P*). Here, *φ*(*P*) is an increasing function of antigen, *P*, that models T cell hyporesponsiveness (Hailemichael et al., 2013). As a simplifying assumption, we do not model the activation of naive T cells explicitly but instead use a saturating Hill function with parameters *k_L_, k_H_* and *X*_0_ chosen so that we obtain biologically realistic behaviours. Furthermore, a key feature of avidity selection is that low levels of antigen expression on DCs stimulate high-avidity T cells and high-levels of antigen expression on DCs stimulate both low- and high-avidity T cells (Alexander-Miller et al., 1996; Bullock et al., 2001; Kedl et al., 2002; Kroger et al., 2008; Rezvani et al., 2011). As such, the activation rate for low-avidity T cells is dependent only on the concentration of DCs with high levels of antigen presentation, *M_H_*, whereas the activation rate for high-avidity T cells is dependent on the total concentration of DCs with low and high levels of antigen presentation, *M_L_ + M_H_*.

### 2.1 Parameter estimates

A list of parameters used in our simulations is given in Table 1. To obtain estimates, we used experimental values for a peptide vaccine against melanomas in humans, but stress that our model readily generalises to other forms of anti-tumour vaccines. Where possible, we have used experimental data from humans to characterise our model parameters; however, specific phenomenological parameters are fit to the results of our previous model.

**Table 1.**
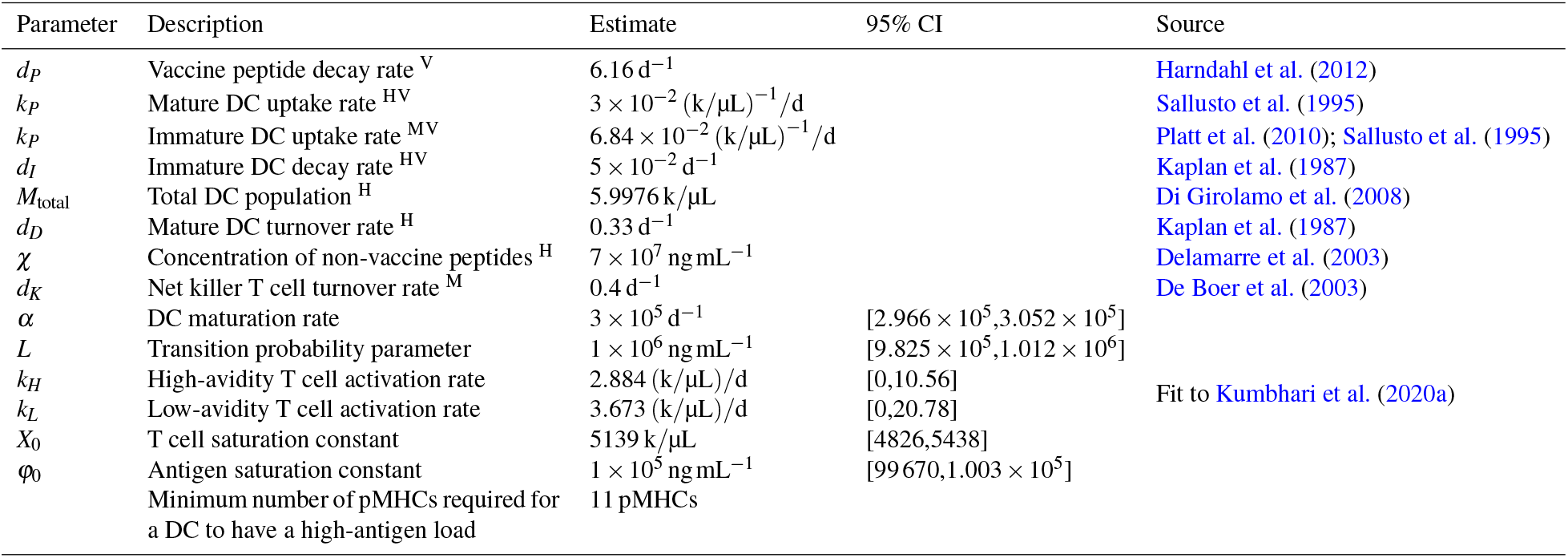
Table of parameters for the ODE model and estimated values. Estimates that are characterised by human, mice and in vitro data are marked with superscripts (·)^H^, (·)^M^, and (·)^V^. Here, d denotes days, and k denotes 10^3^ cells.

#### 2.1.1 Vaccine

In Eq. (1), we assume that the vaccine is given systemically at a fixed dose of *u*_0_ ngmL^−1^ with a dosing interval of *ζ* d equating to a vaccine injection rate of

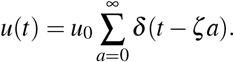

The in vitro decay rate of an immunogenic peptide such as HVDGKILFV is estimated to 6.16 d^-1^, thus, we use a vaccine decay rate, *d_P_*, of 6.16 d^−1^ (Harndahl et al., 2012). We assume that iDCs have an uptake rate of *k_Pi_*, while mature DCs have an uptake rate of *k_P_*. Previously (Kumbhari et al., 2020a), we used human data from the literature (Platt et al., 2010; Sallusto et al., 1995) to estimate a mature DC uptake rate of 3 × 10^−2^ (k/μL)^−1^/d and an immature DC uptake rate of 6.84 × 10^−2^ (k/μL)^−1^/d. Notably, data from Sallusto et al. (1995) shows that while the rate of antigen capture by DCs saturates for large antigen concentrations, the saturation constant associated with this response is large. In other words, even though the rate of antigen capture technically saturates, it *effectively* behaves as a linear function. Thus, as a simplifying assumption, we use mass-action kinetics rather than saturation-type kinetics. Finally, because the vaccine is delivered at *t* = 0, we set *P*(0) = *u*(0) = *u*_0_.

#### 2.1.2 Dendritic cells

In Eq. (2), the rate at which immature DCs turnover, *d_I_*, is 1/20 d^−1^ =5 × 10^−2^ d^−1^ based on human estimates from Kaplan et al. (1987). To calculate the supply rate, *s_I_*, we force the system to be at steady state when there is no antigen, i.e., *P* = 0, equating to *s_I_* – *d_I_I*(0) = 0, or *s_I_* = *d_I_I*(0). The baseline concentration of non-vaccine peptides, *χ*, is 7 × 10^7^ ngmL^−1^ in humans (Delamarre et al., 2003). In Eq. (3) and Eq. (4), the mature DC turnover rate, *d_D_*, is estimated to be 1/72 h^−1^ = 0.33 d^−1^ in humans (Kaplan et al., 1987).

While directly obtaining measurements of DC antigen loads *over time* is challenging, several indirect techniques exists. One such technique involves measuring the percentage of activated low-avidity T cells, which leverages the fact that low-avidity T cell exclusively require high antigen loads for activation. Because the percentage of low-avidity T cells activated (as measured by cytokine secretion and tetramer staining) exhibits a saturation-type response (Bullock et al., 2003), we phenomenologically model the probability of transitioning to a mature DC presenting high levels of surface antigens, *p*_high_(*P*), with a first-order Hill function, i.e.,

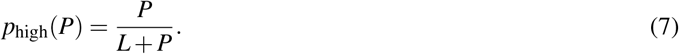

The phenomenological nature of this function means that other sigmoidal functions can be used to model this transition. As the goal here is not to develop a fine-grained model of DC pMHC dynamics (which would be beyond the scope of this paper), we note that a first-order Hill function is sufficiently simple. The model parameter *L*, along with the maturation rate, *α*, is fit to the results of our previous model (Kumbhari et al., 2020a). Details of the fitting procedure are provided in Section 2.2.

For our initial conditions, we note that the total DC population at steady-state conditions, *M*_total_, is reported to be 5.9976k/μL in humans (Di Girolamo et al., 2008). As such, we set *I*(0) = *M*_total_ = 5.9976k/μL. Additionally, we assume that initially there are no mature DCs presenting vaccine-associated peptides, i.e., *M_L,H_* (0) = 0.

#### 2.1.3 T cells

Finally, in Eqs. (5) to (6), both low- and high-avidity T cells decay at rate *d_K_*, which De Boer et al. (2003) estimate to be 0.4 d^−1^ in mice. Motivated by De Boer and Perelson (2013), T cell activation and proliferation is modelled with a saturation function (i.e., a Hill function) with shape parameter *n* = 1. The activation rates *k_H_* and *k_L_*; and saturation constant *X*_0_ are fit to the results of our previous model.

Activation induced cell death (AICD) – also known as “exhaustion”, “senesce”, “adapted” etc. (Blank et al., 2019) – is a phenomenon whereby chronic antigen exposure tempers T cell expansion and is considered a major reason for tumour escape (Hashimoto et al., 2018; June et al., 2018). To model this, we assume our turnover rate, *d_K_*, increases as antigen accumulates. In particular, antigen accumulation, *φ*(*P*), is modelled with the following function:

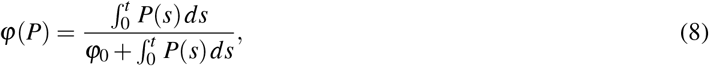

where *φ*_0_ is a saturation constant that is also fitted (details of the fitting procedure are provided in Section 2.2). While the mechanisms behind AICD are unclear (Hashimoto et al., 2018; Blank et al., 2019), it is generally understood that this dysfunctional state occurs due to a history of antigen exposure (Hashimoto et al., 2018). Thus, to account for this history of antigen exposure, we use the integral of *P*, 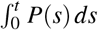, rather than *P* alone. Finally, we assume that initially there are no vaccine-associated effector T cells, i.e., *T_L,H_*(0) = 0.

### 2.2 Parameter fitting

To parametrise our model, we first check for structural identifiability (detailed in Section 2.2.1 below). We then simultaneously fit our model to data generated using our previous model (Kumbhari et al., 2020a), which in turn was based on a model was calibrated to ex vivo human data from Chung et al. (2014), and validated against data from Rezvani et al. (2011) and Hailemichael et al. (2013).

#### 2.2.1 Structurally identifiability analysis

A model is structurally identifiable if, given an infinite amount of noiseless data, all model parameters and initial conditions can be uniquely determined from measurements of its output (Bellman and Astrom, 1970). Moreover, structural identifiability is prerequisite for both prediction (Villaverde et al., 2016; Heinemann and Raue, 2016; Bandara et al., 2009), experimental validation (Villaverde et al., 2016; Walter, 1997; Karr et al., 2015), and importantly practical identifiability (i.e., determining parameter values with noisy data).

To determine if our model is structurally identifiable, we use DAISY (Differential Algebra for Identifiability of SYstems). This software tool checks ODE models with either polynomial or rational nonlinearities for structural identifiability (Bellu et al., 2007). Explicitly, DAISY accepts a set of ODEs describing the state equations (initialised with either known or unknown initial conditions) and uses Ritt’s pseudodivision algorithm to generate an input-output map of the system (i.e., a set of polynomial equations involving only the known variables and their time derivatives). DAISY then uses the Grobner basis of this map to determine if our input-output map is finite-to-one, and thus identifiable (Saccomani and Thomaseth, 2018; Meshkat et al., 2009, 2011, 2012).

A limitation of DAISY is that it only handles rational polynomial nonlinearities and yet Eq. (8) contains an integral. We reconcile this by replacing 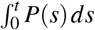 with a dummy variable *P_I_* (defined such that *dP_I_*/*dt* = *P*) and thus leverage the fact that Eqs. (5) and (6) are decoupled from Eqs. (1) to (4). Since we are fitting our model to data generated by our prior model, we assume all state variables are observable.

Using DAISY, we determine that our model is globally structurally identifiable. Our model also implements several first- or second-order Hill functions to model various immunological processes. To assess whether or not structural identifiability depends on the order of the Hill function used, we systematically vary the order from 1 to 10. As a simplifying assumption, we limit these orders to integers. We find that structural identifiability is maintained regardless of which *integer*-order Hill function is used.

#### 2.2.2 Fitting procedure

Structural identifiability establishes that our model can be parametrised via noiseless data. Motivated by this, we calibrate our current model to data from our previous study (Kumbhari et al., 2020a). While our previous model tracked DCs by the number pMHCs being presented, our current model classifies DC antigen loads as being “high” or “low”. To compare the output between the two models, we cluster DC populations as follows. Motivated by reports that as few as four pMHCs suffice to trigger T cell stimulation (Deeg et al., 2013; Varma et al., 2006; Manz et al., 2011), we classify DCs presenting between 1 to 10 pMHCs, i.e., on the same order of magnitude, as having a low antigen density. We then classify DCs presenting over ten pMHCs as having a high antigen load. Our prior work also considered 20 avidity classes, with an avidity state of 1 denoting the lowest and 20 the highest avidity state. Thus, to compare this to our current work, we consider T cells with avidity states ranging from 1–10 as low and states ranging from 11–20 as high.

We then fit our model to a simulated vaccine dose of 7 × 10^5^ ngmL^−1^ given fortnightly. This dosage is chosen as it is similar to the protocols of previous clinical trials (Schwartzentruber et al., 2011; Sosman et al., 2008; Smith et al., 2003; Rezvani et al., 2011). We generate a time trace for the following four variables: DCs with high antigen loads, *M_H_*; DCs with low antigen loads, *M_L_*; high-avidity T cells, *T_H_*; and low-avidity T cells, *T_L_*. Then, for each variable, we calculate the *L*^2^-norm of the error between the time trace predicted by our prior work (after being clustered as per the previous paragraph) and the time trace predicted by our current model. Finally, we use MATLAB’s optimisation routine “fmincon” to find estimates that minimise this aggregate *L*^2^-error. 95% confidence intervals were obtained by bootstrapping residuals 1000 times.

We estimate *α* = 3 × 10^5^d^−1^ (95% CI: [2.966 × 10^5^,3.052 × 10^5^]); *L* = 1 × 10^6^ ngmL^−1^ (95% CI: [9.825 × 10^5^,1.012 × 10^6^]); *k_H_* = 2.884 (k/μL)/d (95% CI: [0,10.56]); *k_L_* = 3.673 (k/μL)/d (95% CI: [0,20.78]); *X*_0_ = 5139k/μL (95% CI: [4826,5438]); and *φ*_0_ = 1 × 10^5^ ngmL^−1^ (95% CI: [99670,1.003 × 10^5^]). This suggests that, relative to the parameters *α, L, X*_0_ and *φ*_0_, *k_L_* and *k_H_* are somewhat poorly identifiable.

As Figure 2 shows, our reduced model underestimates the amplitude of the initial peak for T cells and overestimates the amplitude of secondary T cell peaks. This occurs due to the omission of negative feedback mechanisms such as induced regulatory T cells in our model. Moreover, while other high-low set points could be used, we note that using ten pMHCs provides good *qualitative* agreement with our prior results (see Figure 2).

**Fig. 2.**
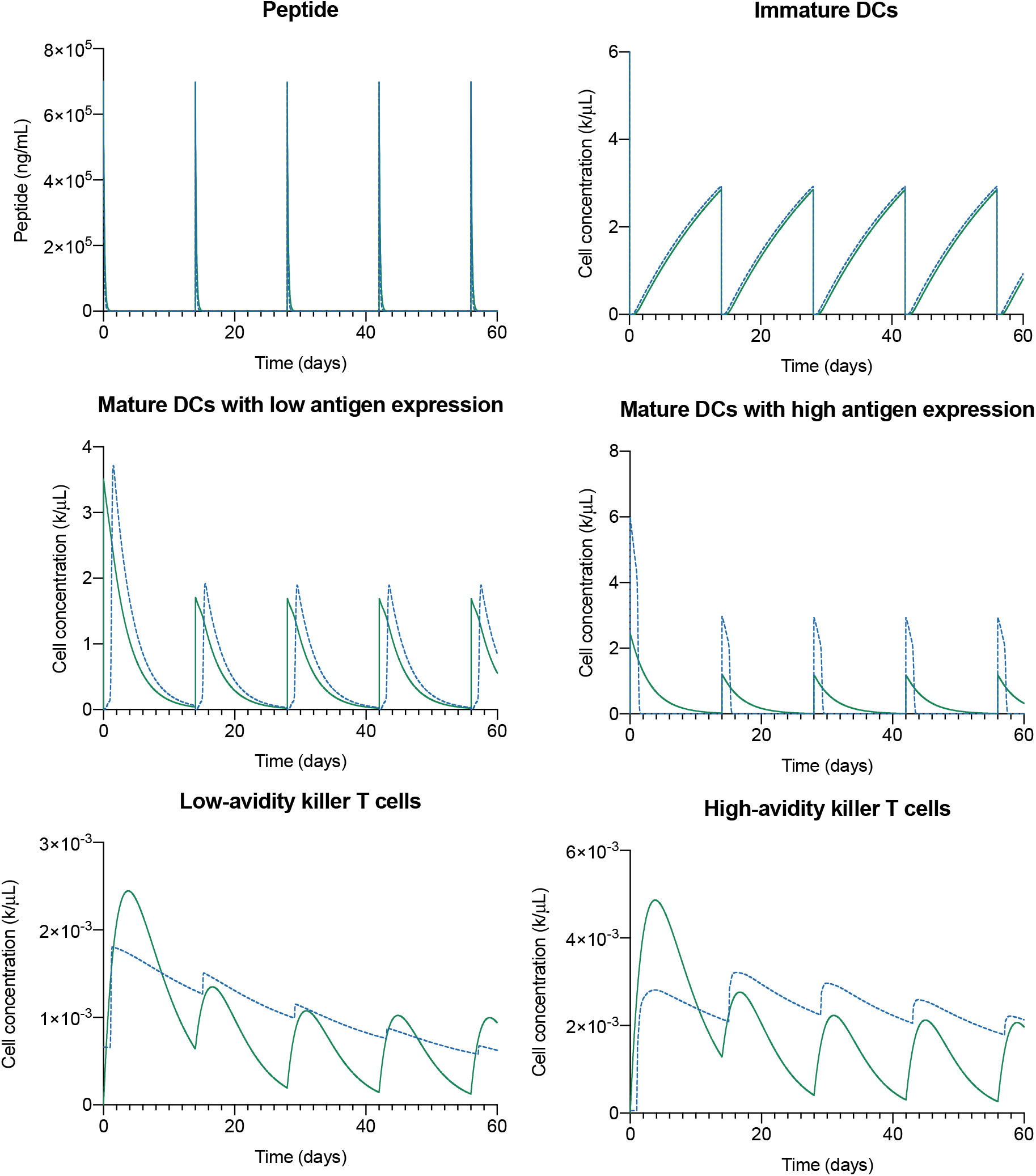
Comparison between the reduced model and our prior work from Kumbhari et al. (2020a) for a dosage of 7 × 10^5^ ngmL^−1^ fortnightly. Here, solid lines correspond to predictions made by our current model and dashed lines to predictions made by our previous model. Simulated cell concentrations are in thousands per micro-litre. Here, we classify a DC presenting between 1 to 10 pMHCs as having a low-antigen load, while anything greater than 10 pMHCs as having a high-antigen load.

## 3 Results

### 3.1 The model is consistent with experimental data

Individualised mathematical models for personalised medicine often necessitate simplicity because of the sparsity of patient data (Andre et al., 2013; Kronik et al., 2010; Gevertz and Wares, 2018). Simple models, however, may be perceived by some to trade mechanistic complexity for abstractions that cannot capture the full scope of experimental data. Here, we validate our reduced model against in vivo murine data from Hailemichael et al. (2013), ex vivo human data from Rezvani et al. (2011) and in vitro data from Wu et al. (2017) and Cawthon et al. (2001).

In Hailemichael et al. (2013), the authors show that repeated vaccination with the gp100 vaccine induces T cell hyporesponsiveness, whereby repeated exposure to an antigen inhibits T cell expansion. To emulate this study, we use a dosage identical to that used in Hailemichael et al. (2013), namely, 100 μg in a 100 μL injection every 42 days, or equivalently 10^6^ ngmL^−1^ every 42 days. Simulating this protocol we find that our model also predicts T cell hyporesponsiveness (see Figure 3A), but the decrease is predicted by our model (20%) is less dramatic than that reported by Hailemichael et al. (2013) (approximately 50%). Since an implicit goal of this study is to develop a minimal model of T cell avidity, we do not include several cell populations (such as myeloid-derived suppressor cells or pro-tumour macrophages) that inhabit the tumour niche and temper T cell expansion. We expect including these factors will produce better agreement with the data from Hailemichael et al. (2013).

**Fig. 3.**
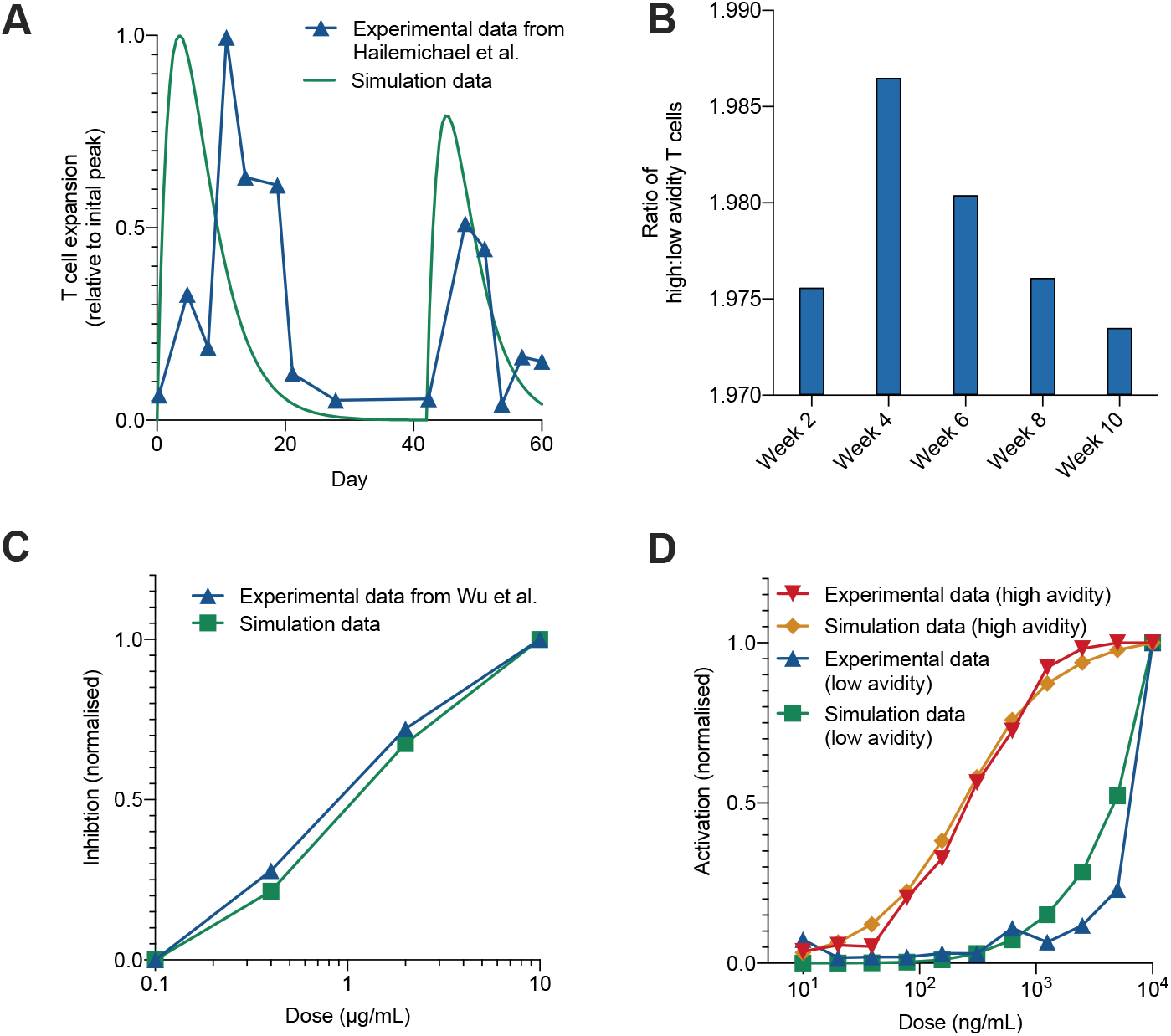
(A) The model predicts T cell hyporesponsiveness as reported by Hailemichael et al. (2013). (B) The model predicts the depletion of high-avidity T cells as reported by Rezvani et al. (2011). (C) Here, we use PD1 expression as an ad-hoc measure of inhibition, which in our model is governed by Eq. (8). PD1 data from Wu et al. (2017) is normalised by mapping the largest MFI to 100% and the lowest MFI to 0%. The average value of Eq. (8), labelled “simulation data”, is similarly normalised. (D) Comparison of activation rates against normalised activation data, quantified via interferon-gamma expression, from Cawthon et al. (2001).

Next, in Rezvani et al. (2011), the authors conduct a small-scale clinical trial with a peptide vaccine, and in doing so observe the depletion of high-avidity T cell (quantified by decreasing ratio of high-avidity to-low-avidity T cell). To test if we also observe a similar depletion in our model, we simulate a dosage of 7 × 10^5^ ngmL^−1^ given every two weeks. We find that after vaccinating at this dosage (see Figure 3B), high-avidity T cells become depleted as observed by Rezvani et al. (2011).

Programmed cell death protein 1 (PD1) is a protein that inhibits T cell activity and is overexpressed on T cells in cancer. In our model, this is implicitly modelled via an increased rate of T-cell turnover (see Eq. (8)). To validate this component of our model, we compare the average value of Eq. (8) against data from Wu et al. (2017) (see Figure 3C), in which the authors show that PD1 expression, quantified via mean fluorescence intensity (MFI), increases with vaccine dosages in vitro (Wu et al., 2017). To simulate Wu et al. (2017)’s in vitro set up, we use a 2-hour dosing frequency and a timespan of 4 days. To simulate the doses reported by Wu et al. (2017), we first note that antigen was distributed across a 24-well plate, which assuming a well working volume of 0.475 mL (Sigma-Aldrich, 2020) and a control volume of 1 mL, implies that 1 μgmL^−1^ of vaccine in vitro equates to a simulated dose of

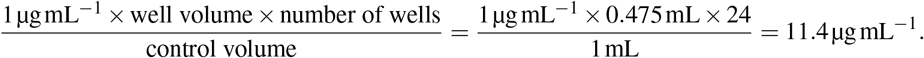

Given that MFI readouts are instrument specific, to compare PD1 MFI readouts, we normalise Wu et al.’s data so that the maximum MFI is mapped to a value of 100%, and the minimum value is mapped to a value of 0%. We then compared this against the average value of Eq. (8), which we similarly normalise. We find that our reduced model agrees well with data from Wu et al. (2017) (see Figure 3C).

Finally, T-cell activation is modelled explicitly via a saturation function (see Eqs. (5) and (6)) and implicitly via a DC pMHC transition probability (see Eqs. (3) and (4)). To validate these components of our model, we compare the average net high- and low-avidity activation rates, *k_H_*(*M_L_* + *M_H_*)/(*X*_0_ + *M_L_* + *M_H_*) and *k_L_M_H_*/(*X*_0_ + *M_H_*), against in vitro T-cell activation data (quantified via interferon-gamma readouts) from Cawthon et al. (2001). To emulate Cawthon et al.’s in vitro set up, we use a 2-hour dosing frequency and a timespan of 1 day. And finally, as data from Cawthon et al. (2001) is normalised to be between 0% and 100%, we similarly normalise our data. We find that our reduced model agrees well with data from Cawthon et al. (2001) (see Figure 3D). Together, these findings show that our model predicts behaviours consistent with the biological literature.

### 3.2 The selection of high-avidity T cells depends synergistically on the schedule rather than the dose or dosing frequency alone

Since our reduced model reproduces key dynamics from the literature, we can leverage our model to identify vaccine schedules that preferentially select for high-avidity T cells. To quantify the selection of high-avidity T cells, we use the *mean avidity difference*

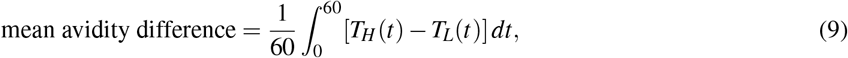

which, unlike the ratio of low-to high-avidity T cells, also accounts for the total T cell concentration. We then perform a global dosage sweep, i.e., simulate combinations of doses ranging from 1 ngmL^−1^ to 1 × 10^9^ ngmL^−1^ with dosing intervals that range from 1 day to 30 days and track the average selection of high-avidity T cells over 60 days (quantified via the mean avidity difference).

We find that a dosage of 1 × 10^3^ngmL^−1^ given every two days maximises the mean avidity difference. A more strategic dosage of 5 × 10^3^ ngmL^−1^ given weekly (see Figure 4) is also comparably effective. Moreover, in Figure 4, for doses between 5 × 10^3^ngmL^−1^ to 1 × 10^4^ngmL^−1^, we notice the formation of a characteristic “ridge”, along which the selection of high-avidity of T cells is robust to the dosing interval. More generally, our simulations suggest that the selection of high-avidity of T cells, as quantified by the avidity difference, is overall more sensitive to dose than to the dosing interval (see Figure 4). This may explain why previous experiments with the gp100 vaccine, which focused primarily on modulating the dosing interval of a high dose vaccine, were unsuccessful in eliciting high-avidity T cells (Schwartzentruber et al., 2011; Sosman et al., 2008; Smith et al., 2003; Rezvani et al., 2011). However, using a low dose alone is also unlikely to induce a significant high-avidity response as T cell expansion is usually proportional to antigen load (Berzofsky et al., 2001). Together, these results suggest that the selection of high-avidity T cells depends synergistically on the dose and the dosing frequency (or schedule), rather than the dosing frequency or dose alone, which is consistent with our previous work.

**Fig. 4.**
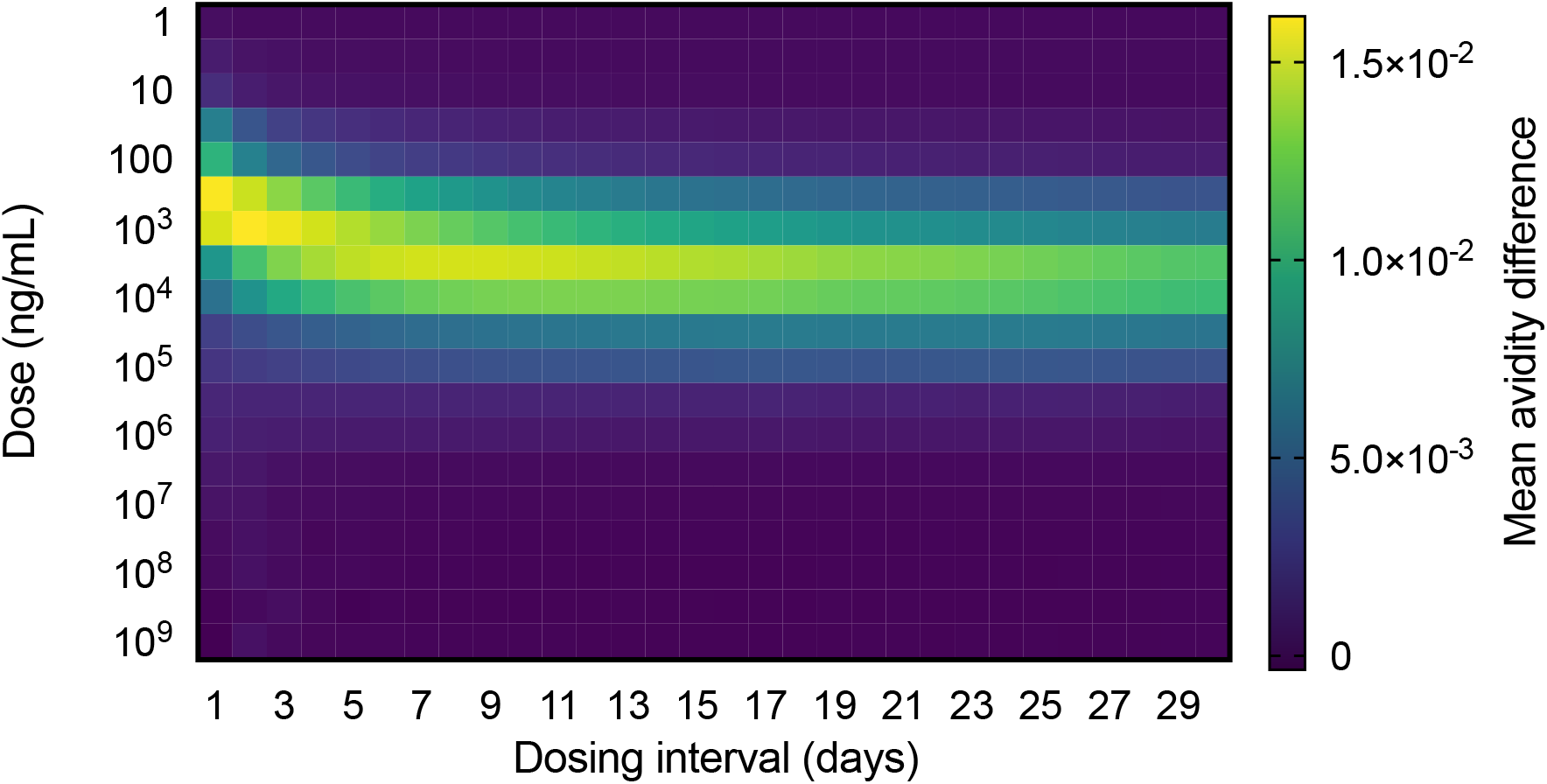
The model predicts a frequent low-dose strategy maximises the selection of high-avidity T cells, which is consistent with prior work. Here, the selection of high-avidity T cells is quantified via the mean-avidity difference, which though *correlated* with tumour clearance is *not* a direct measure of tumour clearance.

### 3.3 Parameter sensitivity changes with dose

In this section, we perform a global sensitivity analysis on several dosages and find that the selection of high-avidity T cells is sensitive to different parameters for different dosages. These sensitivities can be used to eliminate inappropriate (i.e., those that promote low-avidity T cells) dosages. As in Section 2.2, we consider nine dosages, specifically, doses of 10^3^ ngmL^−1^, 7 × 10^5^ ngmL^−1^, or 10^8^ ngmL^−1^; with either weekly, fortnightly, or monthly dosing intervals.

The complexity of individualised immune responses in humans, coupled with a highly heterogeneous tumour microenvironment means that patient data is intrinsically nonlinear (Brodin and Davis, 2017; Zi, 2011). To account for these nonlinear interactions in our sensitivity analysis, we simultaneously vary our parameters over a 100-fold range from their basal values (given in Table 1). Moreover, we generate our samples (*N* = 500) using Latin Hypercube Sampling. To estimate our sensitivity values, we calculate the Spearman Rank Correlation Coefficient (SRCC), *ρ*, between each parameter and the mean avidity difference (defined in Eq. (9)). Table 2 shows the SRCC along with the corresponding *p*-value for several dosages.

**Table 2.** Sensitivity of model parameters for different hypothetical dosages.

| Dosing interval | Dose of $10^3 \text{ ng mL}^{-1}$ | | | Dose of $7 \times 10^5 \text{ ng mL}^{-1}$ | | | Dose of $10^8 \text{ ng mL}^{-1}$ | | |
| --- | --- | --- | --- | --- | --- | --- | --- | --- | --- |
| | Parameter | SRCC | $p$-value | Parameter | SRCC | $p$-value | Parameter | SRCC | $p$-value |
| Weekly | $\alpha$ | 0.4068 | $<10^{-12}$ | $k_H$ | 0.4318 | $<10^{-12}$ | $k_H$ | 0.6452 | $<10^{-12}$ |
| | $k_H$ | 0.3593 | $<10^{-12}$ | $M_{\text{total}}$ | 0.4247 | $<10^{-12}$ | $L$ | 0.2074 | $3.0485 \times 10^{-6}$ |
| | $M_{\text{total}}$ | 0.1357 | 0.0024 | $\alpha$ | 0.1714 | $1.1948 \times 10^{-4}$ | $k_{Pi}$ | 0.0615 | 0.1695 |
| | $k_P$ | 0.0242 | 0.5884 | $\phi_0$ | 0.1263 | 0.0047 | $\chi$ | 0.0090 | 0.8415 |
| | $L$ | 0.0012 | 0.9794 | $d_P$ | 0.1177 | 0.0084 | $M_{\text{total}}$ | 0.0083 | 0.8537 |
| | $k_L$ | -0.0039 | 0.9304 | $L$ | 0.0311 | 0.4872 | $d_P$ | 0.0047 | 0.9170 |
| | $d_K$ | -0.0082 | 0.8543 | $k_P$ | -0.0042 | 0.9248 | $\phi_0$ | -0.0032 | 0.9424 |
| | $\phi_0$ | -0.0385 | 0.3899 | $k_{Pi}$ | -0.0203 | 0.6506 | $d_D$ | -0.0101 | 0.8209 |
| | $d_P$ | -0.0428 | 0.3389 | $d_I$ | -0.0402 | 0.3699 | $X_0$ | -0.0169 | 0.7055 |
| | $d_I$ | -0.0478 | 0.2863 | $k_L$ | -0.0447 | 0.3183 | $d_K$ | -0.0338 | 0.4511 |
| | $k_{Pi}$ | -0.2461 | $2.7460 \times 10^{-8}$ | $d_K$ | -0.1006 | 0.0245 | $k_P$ | -0.0338 | 0.4510 |
| | $d_D$ | -0.3788 | $<10^{-12}$ | $\chi$ | -0.1143 | 0.0106 | $d_I$ | -0.0424 | 0.3434 |
| | $X_0$ | -0.3907 | $<10^{-12}$ | $d_D$ | -0.3933 | $<10^{-12}$ | $\alpha$ | -0.0514 | 0.2508 |
| | $\chi$ | -0.4017 | $<10^{-12}$ | $X_0$ | -0.3994 | $<10^{-12}$ | $k_L$ | -0.5424 | $<10^{-12}$ |
| Fortnightly | $\alpha$ | 0.4070 | $<10^{-12}$ | $k_H$ | 0.4437 | $<10^{-12}$ | $k_H$ | 0.6441 | $<10^{-12}$ |
| | $k_H$ | 0.3592 | $<10^{-12}$ | $M_{\text{total}}$ | 0.4064 | $<10^{-12}$ | $L$ | 0.2074 | $3.0610 \times 10^{-6}$ |
| | $M_{\text{total}}$ | 0.1357 | 0.0024 | $\alpha$ | 0.1819 | $4.3965 \times 10^{-5}$ | $k_{Pi}$ | 0.0613 | 0.1707 |
| | $k_P$ | 0.0244 | 0.5859 | $d_P$ | 0.1047 | 0.0193 | $\chi$ | 0.0090 | 0.8413 |
| | $L$ | 0.0013 | 0.9776 | $\phi_0$ | 0.0982 | 0.0282 | $M_{\text{total}}$ | 0.0071 | 0.8737 |
| | $k_L$ | -0.0040 | 0.9293 | $L$ | 0.0266 | 0.5522 | $d_P$ | 0.0070 | 0.8762 |
| | $d_K$ | -0.0080 | 0.8580 | $k_P$ | -0.0126 | 0.7789 | $\phi_0$ | -0.0015 | 0.9737 |
| | $\phi_0$ | -0.0392 | 0.3817 | $k_{Pi}$ | -0.0312 | 0.4869 | $d_D$ | -0.0095 | 0.8316 |
| | $d_P$ | -0.0429 | 0.3386 | $k_L$ | -0.0484 | 0.2803 | $X_0$ | -0.0173 | 0.6988 |
| | $d_I$ | -0.0475 | 0.2888 | $d_I$ | -0.0581 | 0.1943 | $d_K$ | -0.0326 | 0.4665 |
| | $k_{Pi}$ | -0.2464 | $2.6398 \times 10^{-8}$ | $d_K$ | -0.0731 | 0.1026 | $k_P$ | -0.0353 | 0.4311 |
| | $d_D$ | -0.3787 | $<10^{-12}$ | $\chi$ | -0.1183 | 0.0081 | $d_I$ | -0.0432 | 0.3351 |
| | $X_0$ | -0.3906 | $<10^{-12}$ | $d_D$ | -0.4031 | $<10^{-12}$ | $\alpha$ | -0.0503 | 0.2619 |
| | $\chi$ | -0.4015 | $<10^{-12}$ | $X_0$ | -0.4097 | $<10^{-12}$ | $k_L$ | -0.5418 | $<10^{-12}$ |
| Monthly | $\alpha$ | 0.4069 | $<10^{-12}$ | $k_H$ | 0.4495 | $<10^{-12}$ | $k_H$ | 0.6430 | $<10^{-12}$ |
| | $k_H$ | 0.3591 | $<10^{-12}$ | $M_{\text{total}}$ | 0.3952 | $<10^{-12}$ | $L$ | 0.2070 | $3.1920 \times 10^{-6}$ |
| | $M_{\text{total}}$ | 0.1356 | 0.0024 | $\alpha$ | 0.1868 | $2.7250 \times 10^{-5}$ | $k_{Pi}$ | 0.0617 | 0.1684 |
| | $k_P$ | 0.0244 | 0.5867 | $d_P$ | 0.0953 | 0.0332 | $\chi$ | 0.0096 | 0.8306 |
| | $L$ | 0.0010 | 0.9819 | $\phi_0$ | 0.0823 | 0.0658 | $d_P$ | 0.0091 | 0.8383 |
| | $k_L$ | -0.0040 | 0.9297 | $L$ | 0.0239 | 0.5930 | $M_{\text{total}}$ | 0.0063 | 0.8877 |
| | $d_K$ | -0.0080 | 0.8588 | $k_P$ | -0.0181 | 0.6870 | $\phi_0$ | $-8.2176 \times 10^{-5}$ | 0.9985 |
| | $\phi_0$ | -0.0393 | 0.3798 | $k_{Pi}$ | -0.0369 | 0.4097 | $d_D$ | -0.0092 | 0.8376 |
| | $d_P$ | -0.0428 | 0.3394 | $k_L$ | -0.0496 | 0.2684 | $X_0$ | -0.0174 | 0.6977 |
| | $d_I$ | -0.0475 | 0.2889 | $d_K$ | -0.0586 | 0.1911 | $d_K$ | -0.0312 | 0.4866 |
| | $k_{Pi}$ | -0.2464 | $2.6587 \times 10^{-8}$ | $d_I$ | -0.0681 | 0.1281 | $k_P$ | -0.0363 | 0.4178 |
| | $d_D$ | -0.3789 | $<10^{-12}$ | $\chi$ | -0.1225 | 0.0061 | $d_I$ | -0.0423 | 0.3452 |
| | $X_0$ | -0.3906 | $<10^{-12}$ | $d_D$ | -0.4100 | $<10^{-12}$ | $\alpha$ | -0.0511 | 0.2543 |
| | $\chi$ | -0.4015 | $<10^{-12}$ | $X_0$ | -0.4134 | $<10^{-12}$ | $k_L$ | -0.5412 | $<10^{-12}$ |

We find that for large doses, the sensitivity of DC maturation rate to antigen, *L*, and the total DC concentration, *M*_total_ are parameters that are positively correlated with the promotion of high-avidity T cells. By contrast, the DC antigen consumption rates, *k_P_* and *k_Pi_*, and the T cell saturation constant (a measure of how sensitivity T cell activation is to mature DC concentrations), *X*_0_, are parameters that are negatively correlated with the promotion of high-avidity T cells, i.e., they are positively correlated with the selection of low-avidity T cells. This suggests that the presentation of antigen on mature DCs may be driving the selection of high-avidity T cells in our model, which is consistent with the literature (Gerner et al., 2017; van Stipdonk et al., 2001). We also find that as the vaccine dose increases, the sensitivity of *α*, the DC maturation rate decreases in our simulations. Again, this suggests that both antigen presentation and DC activation dynamics drive the promotion of high-avidity T cells.

By estimating the parameter sensitivity of these candidate dosages, we can use a patient’s history and clinical presentation to screen for suitable dosages. For example, a common co-morbidity for melanoma patients is diabetes (Lee et al., 2015). Diabetes is known to decrease phagocytosis by immune cells (Geerlings and Hoepelman, 1999), which in our model would correspond to a decreased rates of antigen uptake, *k_P_* and *k_Pi_*. The goal here is to identify dosages where the selection of high-avidity T cells is negatively correlated with both *k_P_* and *k_Pi_*. Referring to Table 2, we identify 7 × 10^5^ ngmL^−1^ given either weekly, fortnightly or monthly as suitable candidate dosages.

As an additional example, consider a patient presenting with a history of heart disease another common co-morbidity in skin cancer patients. Coronary artery disease results in a have a lower number of circulating DCs in patients (Van Vre et al., 2011). In our model, this corresponds to a decreased DC concentration, *M*_total_. The goal here is to identify dosages where the selection of high-avidity T cells is weakly correlated with *M*_total_. Referring to Table 2, we identify 1 × 10^8^ ngmL^−1^ given either weekly, fortnightly or monthly possible candidate dosages. Together, these examples illustrate how our model can be leveraged to personalise dosages based on a patient’s history and other conditions.

## 4 Discussion

Personalising treatment schedules to induce high-avidity T cells is a promising new approach to maintaining immunity against certain cancers such as melanoma. Here we develop a simple ODE model of T cell avidity that is validated against several experimental datasets. We then use our model to suggest therapy schedules based on a table of sensitivities. This method involves first using a patient’s history and clinical presentation to determine which parameters are expected to have changed, and then referring to a table of parameter sensitivities to eliminate inappropriate vaccine schedules. Importantly, our study is a proof-of-concept study and still requires experimental validation.

Since this study aims to develop a minimal model of avidity selection, our model makes several simplifying biological assumptions. For example, we do not account for certain immunological processes such as the induction of regulatory T cells, lymphocyte trafficking, and cytokine secretion. These processes were, however, modelled in our prior work, against which we calibrated our model. Additionally, we only considered two avidity states, low or high, despite avidity likely existing on a continuous spectrum. Since most experimental studies only report on low- and high-avidity populations, using a system of ODEs (rather than a similar system of PDEs) makes the model more amenable to experimental validation.

As an additional simplification, we assumed the probability of an immature DC transitioning to a mature DC presenting high levels of surface antigens, *p*_high_(*P*), was dependent only on the concentration of antigen. Biologically, this probability depends on additional, more dynamic factors, such as co-signalling pathways (Chen and Flies, 2013). Importantly, our model is not specific to peptide vaccines, and we expect that our model can also apply to newer, neo-antigen-based T cell vaccines.

As an alternative to parametrising and then optimising a model to patient data (which may be difficult), we propose using a table of sensitivities to screen to suitable dosages. This table of sensitivities involves performing a sensitivity analysis on model parameters for a set of different simulated dosages. Of these dosages, some, when clinically trialled, were found to promote the low-avidity T cells over high-avidity T cells (Hailemichael et al., 2013; Rezvani et al., 2011; Schwartzentruber et al., 2011). Nonetheless, we included these dosages as we could not rule out the possibility that these dosages are optimised to elicit anti-tumour immunity by additional, non-avidity-based mechanisms not considered in our model. Indeed, we argue that, under the right conditions (i.e., those identified via our table of sensitivities), these dosages may enhance the selection of high-avidity T cells. To ensure our schedules are also practical, we limit our simulated dosing intervals to weekly, fortnightly or monthly intervals. Consequently, we did not include the optimal dosage of 1 × 10^3^ngmL^−1^ given every two days in our table of candidate schedules. Moreover, while these dosing intervals were chosen for their practicality, an alternative approach not explored in this study involves using control theory to identify an optimal vaccine strategy. Indeed, using a control-theoretic approach may identify dosages that maximise the selection of high-avidity T cells beyond what we identified. Finally, while our findings still require preclinical validation, we anticipate that our simulated dosages are safe. This is based on studies in which limited adverse side effects are reported (Hailemichael et al., 2013; Rezvani et al., 2011; Schwartzentruber et al., 2011). As such, we predict that under the right circumstances (such as those suggested by Table 2), these dosages can safely elicit high-avidity T cells.

Notably, the schedules identified here all aim to maximise the mean avidity difference over 60 days. We used the avidity difference rather than the ratio of high-to low-avidity T cells, as it not only penalises the selection of low-avidity T cells but also accounts for the total concentration of T cells induced, which is an important predictor of treatment efficacy (Kittlesen et al., 1998). However, as a metric for the selection of high-avidity T cells, the mean avidity difference has limitations. For example, optimising the mean avidity difference results in all-or-nothing control, whereby a response that elicits a low total T cell count has a higher payoff than one that promotes low-avidity T cells at a higher concentration. While this allows us to account implicitly for the inhibition of high-avidity T cells by low-avidity T cells (Chung et al., 2014), this also results in parameter sensitivities that suggest T cell hyporesponsiveness is preferable over the stimulation of low-avidity T cells. This could be addressed by using a metric that penalises both the selection of low-avidity T cells and low total T cell concentrations. Vaccine protocols also need to account for factors other than T cell avidity, such as toxicity constraints and off-target reactions (Tigue et al., 2007), which are factors that our model does not include. Developing a selection metric that accounts for these factors will be the subject of future investigations.

Overall, our findings still require substantial experimental validation, which is a priority for future work. Nonetheless, they provide a vital proof-of-concept link between a phenotypic model of avidity selection and identifying and eliminating sub-therapeutic vaccine schedules, which may help in inducing durable anti-tumour immunity.

## Acknowledgements

This work was supported by an Australian Government Research Training Program Scholarship (AK); Australian Research Council Discovery Project [DP180101512] (PSK and DR); and by the US Department of Defense Breast Cancer Research Program [W81XWH-11–1–0548] (PPL).

